# Influence of Strain Post-Processing on Brain Injury Prediction

**DOI:** 10.1101/2021.07.15.452485

**Authors:** Madelen Fahlstedt, Shiyang Meng, Svein Kleiven

## Abstract

Finite element head models are a tool to better understand brain injury mechanisms. Many of the models use strain as output but with different percentile values such as 100^th^, 95^th^, 90^th^, and 50^th^ percentiles. Some use the element value, whereas other use the nodal average value for the element. Little is known how strain post-processing is affecting the injury predictions and evaluation of different prevention systems. The objective of this study was to evaluate the influence of strain output on injury prediction and ranking.

Two models with different mesh densities were evaluated (KTH Royal Institute of Technology head model and the Total Human Models for Safety (THUMS)). Pulses from reconstructions of American football impacts with and without a diagnosis of mild traumatic brain injury were applied to the models. The value for 100^th^, 99^th^, 95^th^, 90^th^, and 50^th^ percentile for element and nodal averaged element strain was evaluated based on peak values, injury risk functions, injury predictability, correlation in ranking, and linear correlation.

The injury risk functions were affected by the post-processing of the strain, especially the 100^th^ percentile element value stood out. Meanwhile, the area under the curve (AUC) value was less affected, as well as the correlation in ranking (Kendall’s tau 0.71-1.00) and the linear correlation (Pearson’s r^2^ 0.72-1.00). With the results presented in this study, it is important to stress that the same post-processed strain should be used for injury predictions as the one used to develop the risk function.

## Introduction

Head injuries are a significant problem in society that can cause both acute and long-term consequences. Mild traumatic brain injuries (mTBIs) are a growing concern in society among all age groups. It is estimated that 5.3 million sustain an mTBI every year in the US (Roozenbeek et al., 2013). There are efforts to decrease the number of mTBIs, for example, by trying to improve the prevention strategies through a better understanding of the injury mechanisms. A tool that is more and more commonly used is finite element (FE) head models. More than a dozen FE head models developed worldwide are used in different situations related to injury prevention. The models have different designs, such as numbers of elements, contacts, material models, geometry, etc. (Fahlstedt et al., 2021; Giudice et al., 2019).

In most cases, the peak strain is used as an injury predictor for the models. However, there is some difference in how the peak values are presented. For some models, the 100^th^ percentile value of the maximum principal strain (MPS) is used (Kleiven, 2007; Trotta et al., 2020), whereas others use the 95^th^ percentile value (Elkin et al., 2019; Wu et al., 2020), the 90^th^ percentile value (Ghajari et al., 2017), or the 50^th^ percentile value (Gabler et al., 2018). The motivation to use lower percentile values has been to eliminate numerical instabilities (Gabler et al., 2016; Hajiaghamemar et al., 2020; Panzer et al., 2012).

Another difference has been that some have presented the element strain based on nodal averaging (e.g., Kleiven (2007)), and other studies have shown the strain based directly on the element value (e.g. (Hernandez et al., 2019)). Little or no motivation is given why the nodal averaged element value or direct element value is used. There is also limited knowledge of how this is affecting the results.

For several of the developed models, injury risk functions have been developed for mTBI. These injury risk functions have been used to evaluate, compare and rate different prevention systems (e.g. Clark et al., 2018; Elkin et al., 2019; Fahlstedt et al., 2021). The objective of this study was to evaluate the influence of strain post-processing on injury prediction and ranking by using two different FE models with different mesh densities.

## Methods

The KTH Royal Institute of Technology model (Kleiven, 2007) and the Total Human Body Model for Safety (THUMS) (Atsumi et al., 2016) were used in this study. A summary of the models is found in Table A1 and Table A2.

The dataset of impacts in American football presented by Sanchez et al. (2019) was used. The dataset includes 53 cases (20 with mTBI and 33 without mTBI). The linear acceleration and angular velocity were obtained from reconstructions with anthropometric test devices (Newman et al., 1999, 2005; Pellman et al., 2003; Sanchez et al., 2019). The linear acceleration and angular velocity for the head’s x-, y-, and z-axes were applied to the center of gravity of the head models. The time window for the different pulses suggested by Sanchez et al. (2019) was used. For both models, the simulations were run on a local Linux cluster with four processors. For the KTH model, LS Dyna revision 12.0 was used, and for THUMS revision 9.1, both with double precision.

The results from the simulations were analyzed in LS PrePost (version 4.3.7). The output from the models was the first principal Green-Lagrange strain, which was calculated with either nodal averaged element or element value. Nodal averaging is used to average the element strain values over nodes common to two or more elements. This is used to obtain continuous strain fields over element boundaries since the direct element values otherwise are constant over each element and discontinuous between elements. The values were presented for the 100^th^, 99^th^, 95^th^, 90^th^, and 50^th^ percentile values for maximum value in each element for the whole brain and the brainstem.

Injury risk curves with survival analysis based on the Weibull distribution were developed. In the survival analysis, the non-concussion cases were treated as right-censored and the concussion cases as left-censored.

Based on the injury risk functions, the receiver operating characteristic (ROC) curve was generated. The correlation of ranking was evaluated with Kendall’s tau and linear correlation with Pearson’s coefficient of determination (r^2^). All the statistical analysis was performed in Matlab (version 2019a, The MathWorks, Inc., Natick, Massachusetts, United States).

## Results

The THUMS model showed a decreasing difference of the mean value of the 53 cases between nodal averaged element and element value with decreased percentile value (Figure 1). A smaller difference among the high percentile values between the element value and the nodal averaged element value was seen for the brainstem of THUMS (Figure 1b). The same trend was seen for the KTH model (Figure 2). The injury risk functions for the two models are shown in Figure 3 and Figure 4. The constants for the survival analysis function for both models are presented in Table A3.

**Figure 1.**
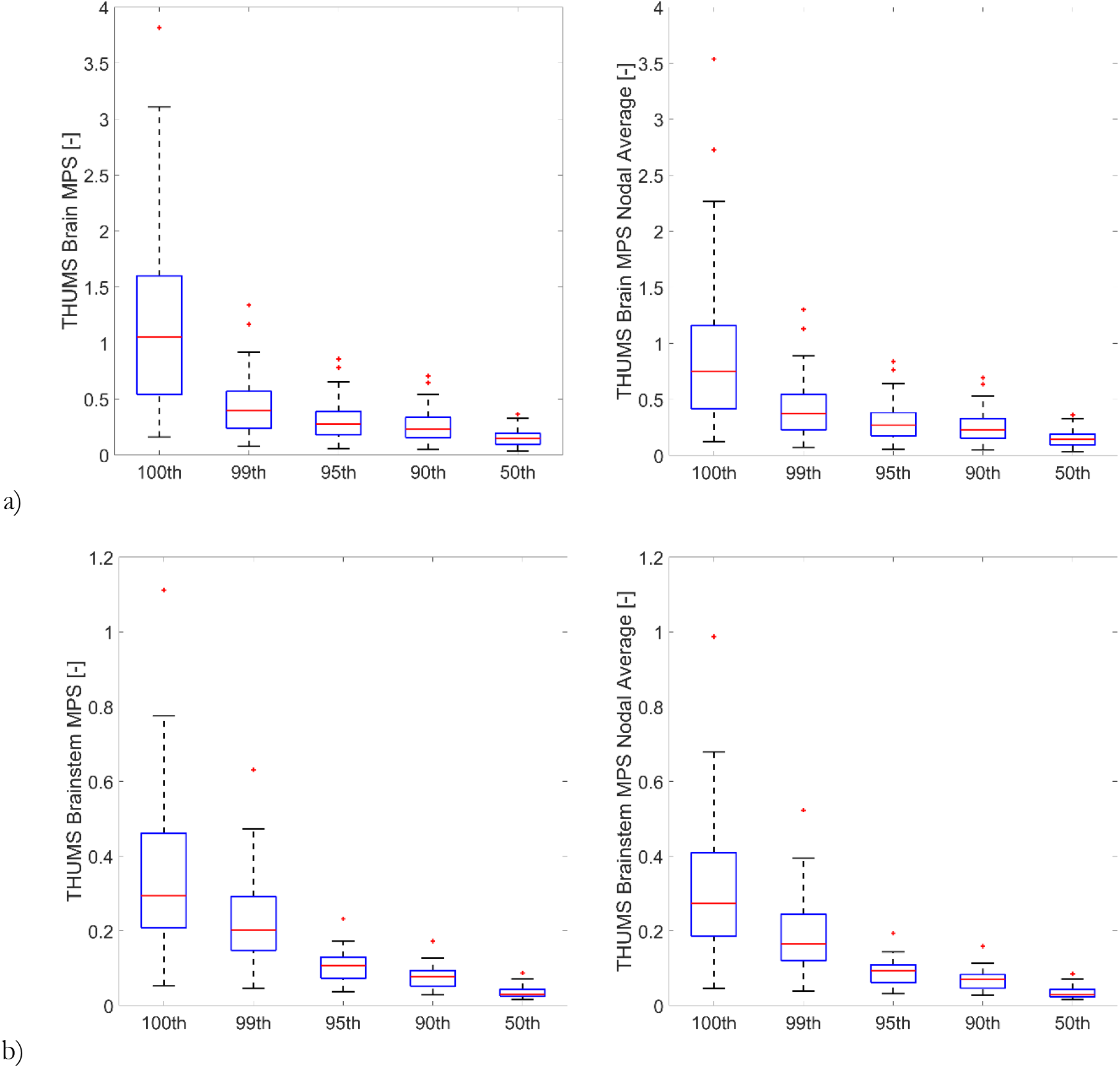
Boxplot of MPS of THUMS for element (left) and nodal averaged element (right) value and the different percentile values (100^th^, 99^th^, 95^th^, 90^th^, and 50^th^) for a) brain, b) brainstem

**Figure 2.**
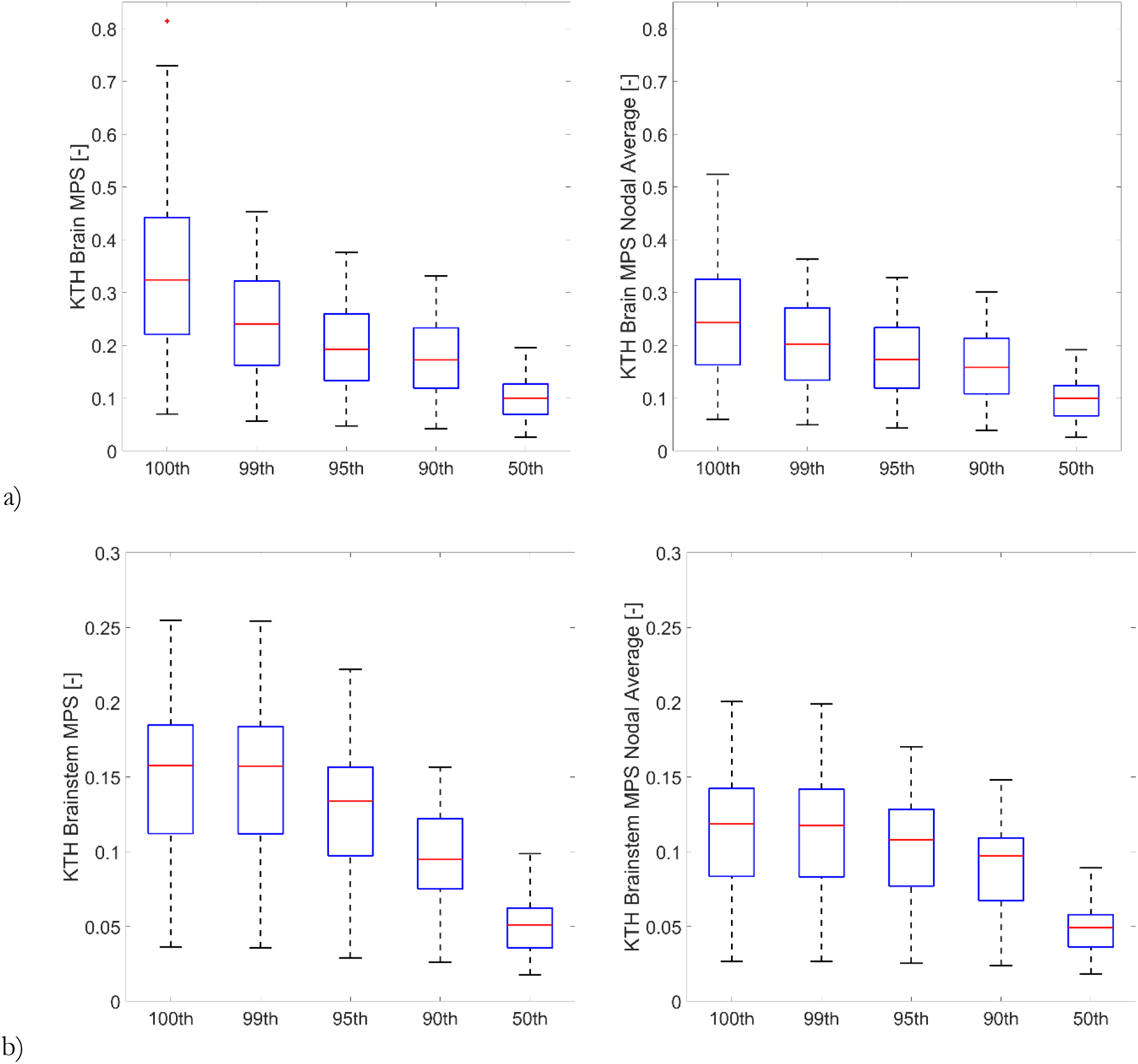
Boxplot of MPS of KTH for element (left) and nodal averaged element (right) value and the different percentile values (100^th^, 99^th^, 95^th^, 90^th^, and 50^th^) for a) brain, b) brainstem

**Figure 3.**
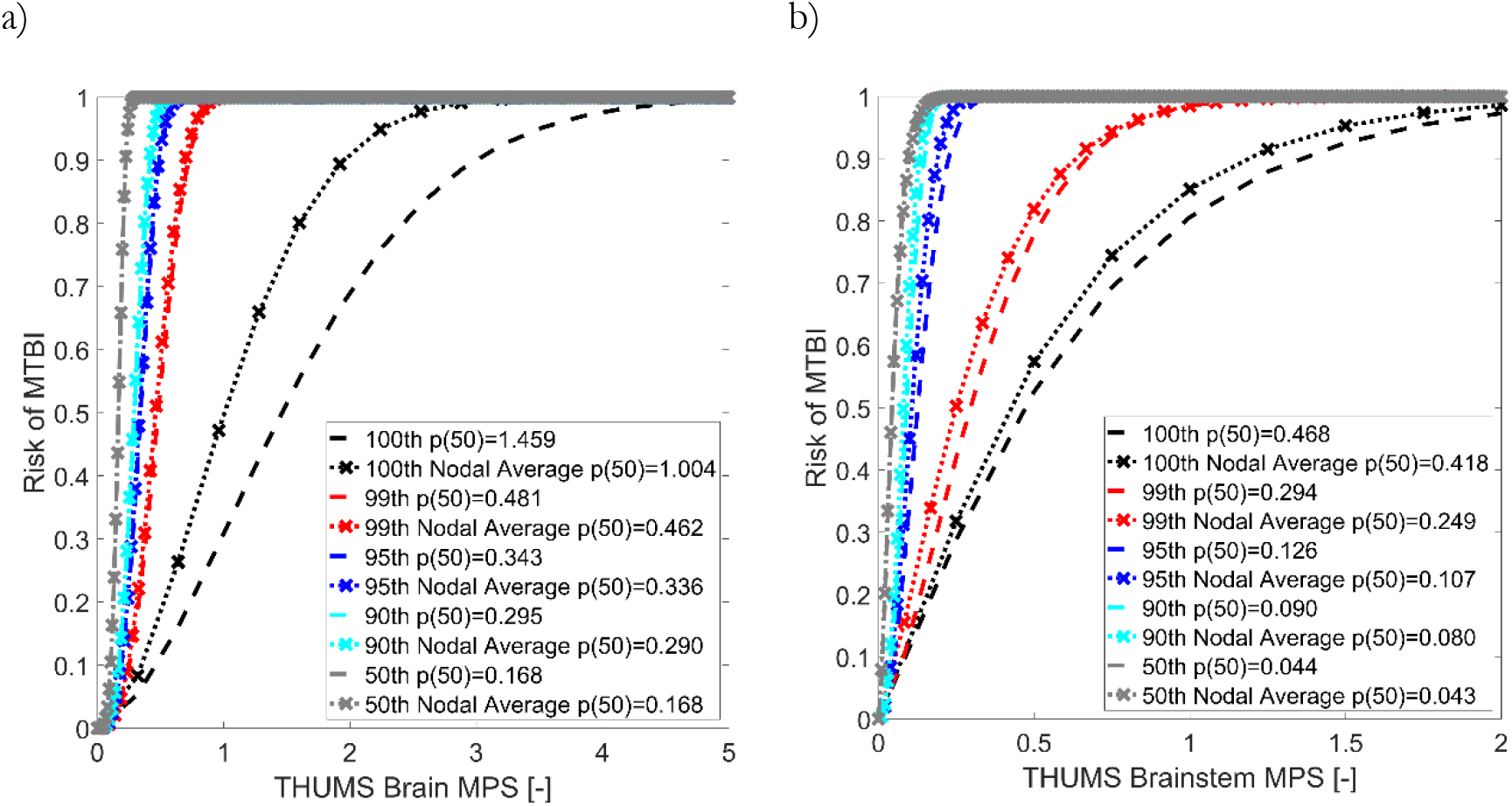
Risk curves for THUMS of MPS for with and without nodal averaged element and the different percentile values (100^th^, 99^th^, 95^th^, 90^th^, and 50^th^) for a) brain, b) brainstem

**Figure 4.**
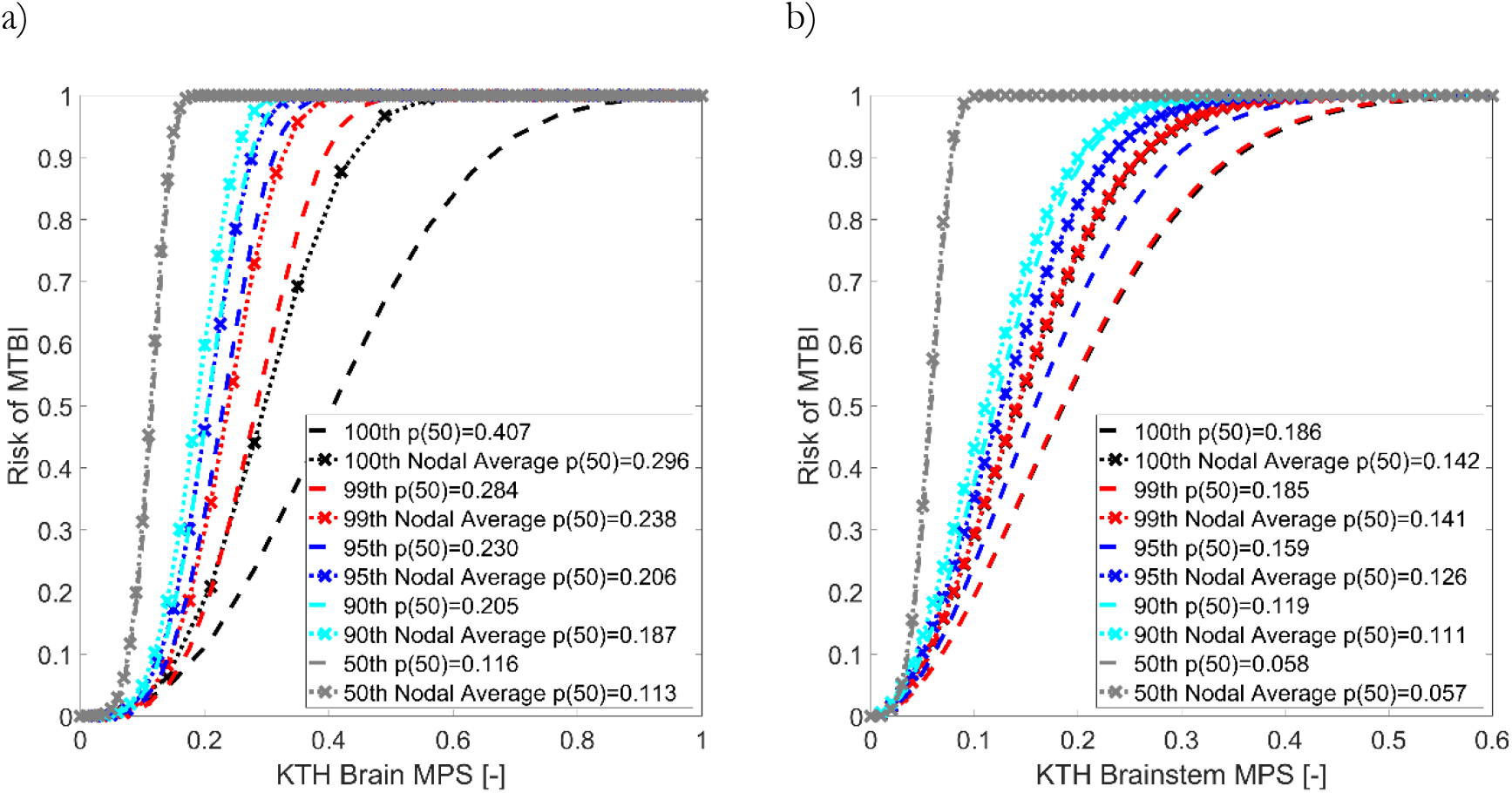
Risk curves for the KTH model of MPS for with and without nodal averaged element and the different percentile values (100^th^, 99^th^, 95^th^, 90^th^, and 50^th^) for a) brain, b) brainstem

The highest AUC value for the THUMS model for the whole brain was found for nodal averaged 50^th^ percentile value for both nodal averaged element and element value but with small variations (Figure 5a left). For the brainstem, the lowest AUC value for both element and the nodal averaged element value was found for the 50^th^ percentile (Figure 5a right).

**Figure 5.**
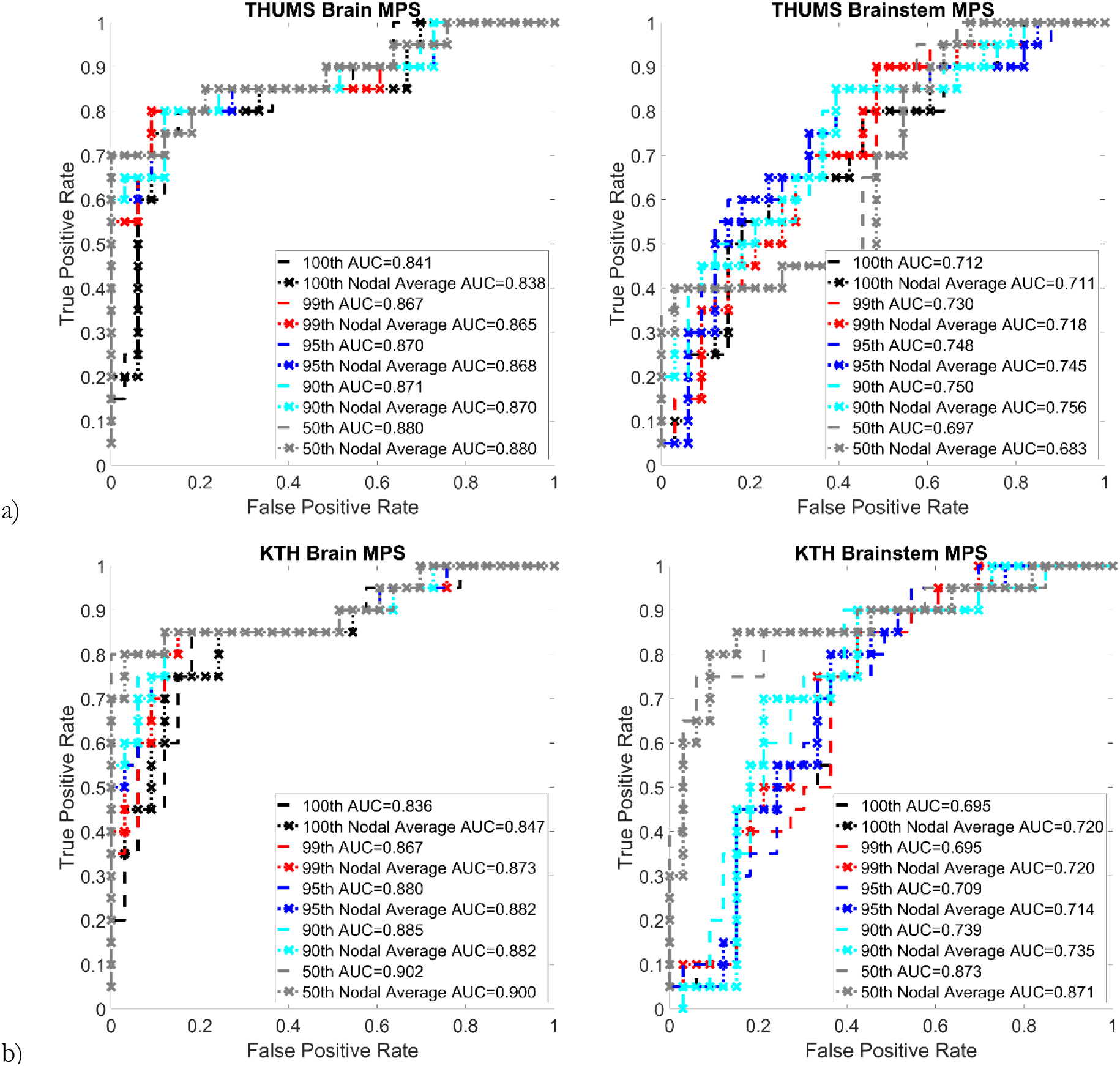
ROC curves (first row THUMS and second row KTH, left column whole brain, right column brainstem).

For the KTH model, the highest AUC value was also found for the 50^th^ percentile value independent of how the strain was post-processed. However, the brainstem had a lower AUC value compared to the whole brain (Figure 5).

An illustration of the strain over time and the peak value for each element together with the different percentile values can be seen in Figure 6 and Figure 7. The case with the highest peak strain and lowest peak strain for both models were chosen as examples.

**Figure 6.**
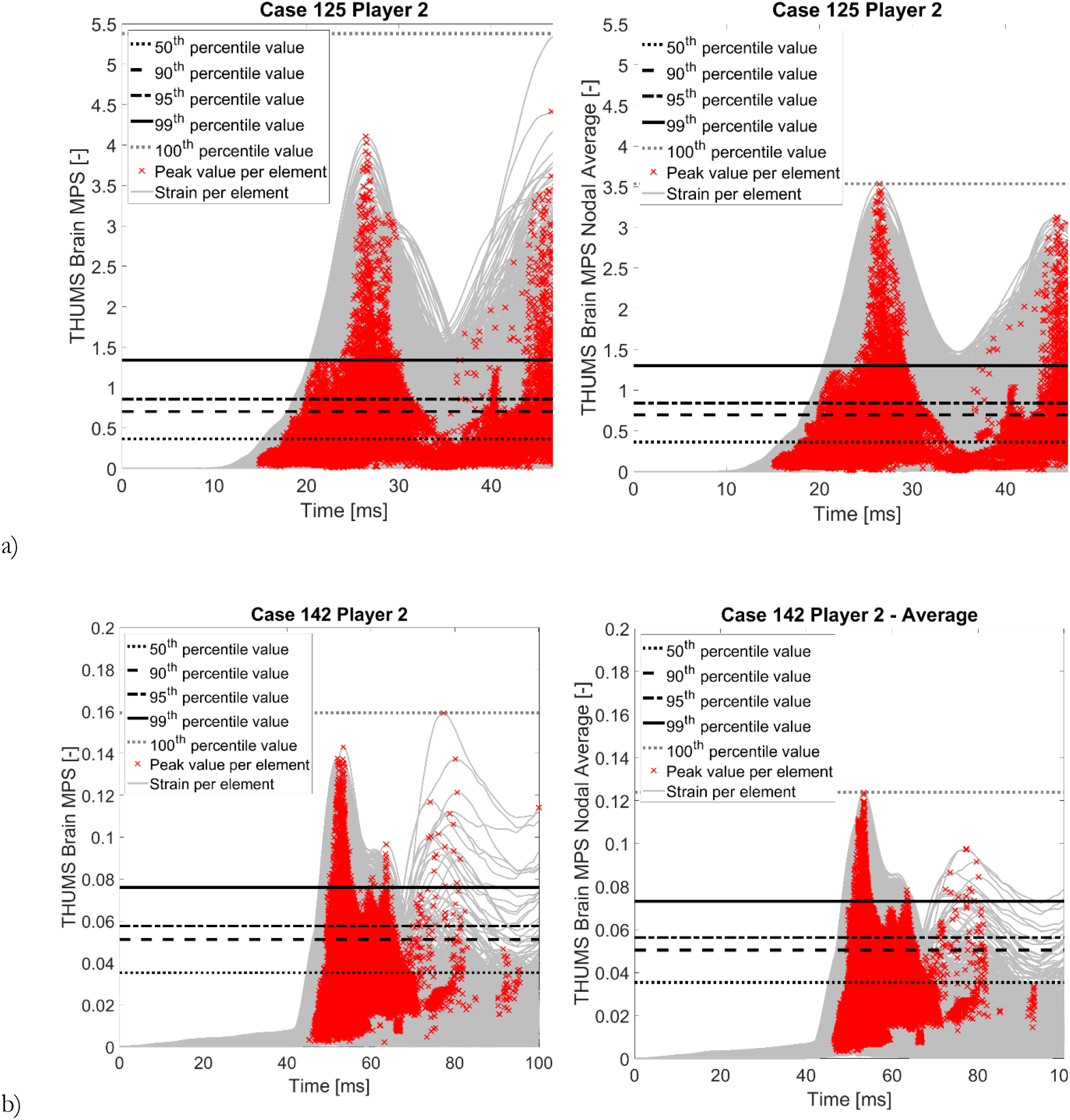
Strain over time for the THUMS model for the brain tissue (grey lines), where the peak value for each element is marked with a red cross (x) and the horizontal lines shows the 50^th^ to 100^th^ percentile values; a) the case with highest peak strain for the THUMS model, b) the case with lowest peak strain for the THUMS model.

**Figure 7.**
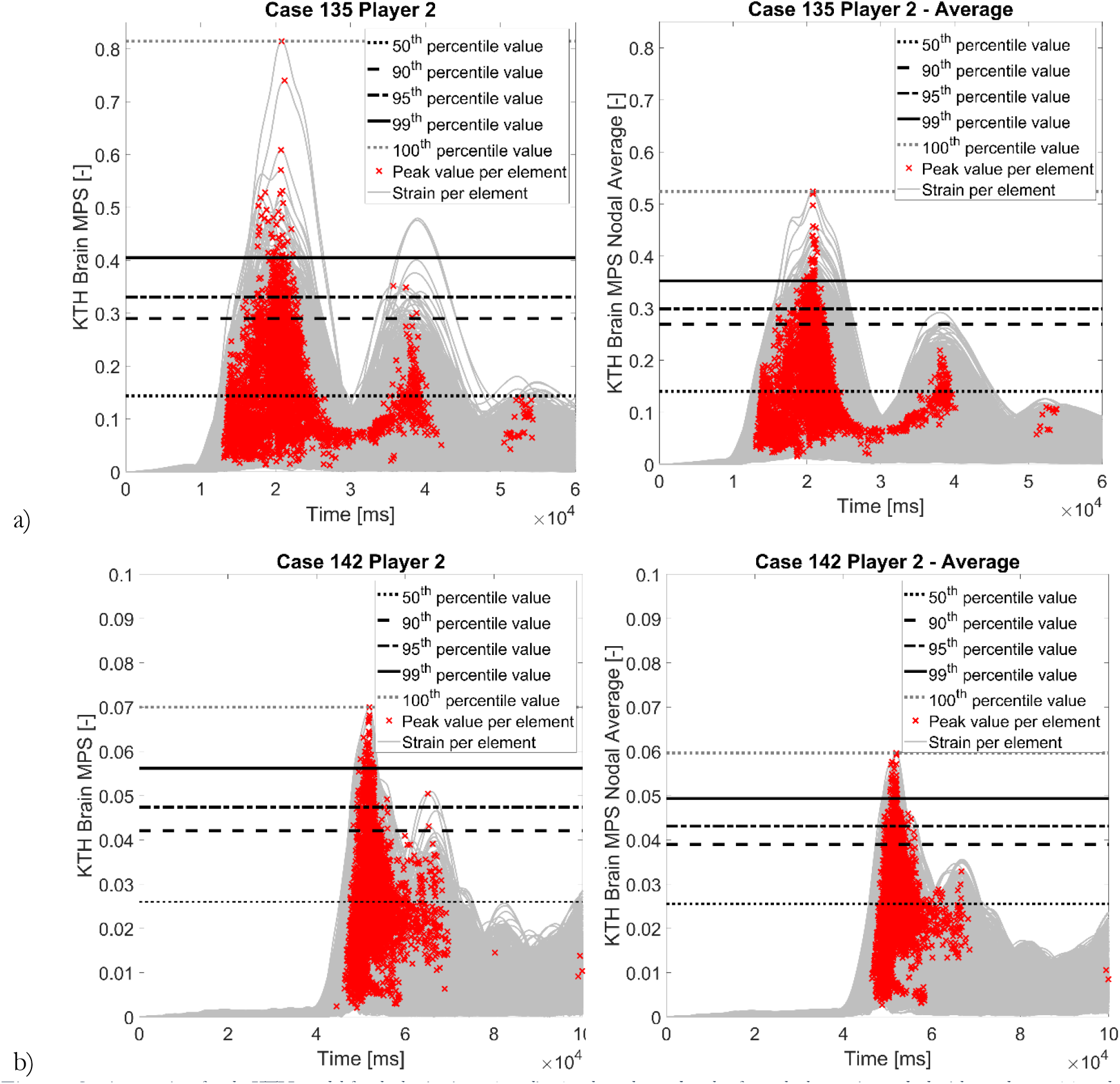
Strain over time for the KTH model for the brain tissue (grey lines), where the peak value for each element is marked with a red cross (x) and the horizontal lines shows the 50^th^ to 100^th^ percentile; a) the case with highest peak strain for the KTH model, b) the case with lowest peak strain for the KTH model.

The correlation in ranking and linear correlation shown in Table 1 for both models. The scatter plots are presented in the Appendix.

**Table 1.** Correlation of ranking (Kendall’s Tau) and linear correlation (Pearson’s coefficient of determination r^2^) for element and nodal averaged element strain and for the different percentile values.

| | | Kendall's Tau | | Pearson's $r^2$ | |
| --- | --- | --- | --- | --- | --- |
|  |  | THUMS | KTH | THUMS | KTH |
| Nodal Averaged Element Strain | 100<sup>th</sup>- 99<sup>th</sup> | 0.84 | 0.91 | 0.92 | 0.95 |
|  | 100<sup>th</sup>- 95<sup>th</sup> | 0.82 | 0.89 | 0.88 | 0.93 |
|  | 100<sup>th</sup>- 90<sup>th</sup> | 0.82 | 0.88 | 0.86 | 0.92 |
|  | 100<sup>th</sup>- 50<sup>th</sup> | 0.74 | 0.78 | 0.79 | 0.82 |
|  | 99<sup>th</sup>- 95<sup>th</sup> | 0.96 | 0.97 | 0.99 | 0.99 |
|  | 99<sup>th</sup>- 90<sup>th</sup> | 0.95 | 0.96 | 0.98 | 0.99 |
|  | 99<sup>th</sup>- 50<sup>th</sup> | 0.87 | 0.86 | 0.94 | 0.93 |
|  | 95<sup>th</sup>- 90<sup>th</sup> | 0.99 | 0.98 | 1.00 | 1.00 |
|  | 95<sup>th</sup>- 50<sup>th</sup> | 0.89 | 0.88 | 0.96 | 0.96 |
|  | 90<sup>th</sup>- 50<sup>th</sup> | 0.90 | 0.89 | 0.97 | 0.97 |
| Element Strain | 100<sup>th</sup>- 99<sup>th</sup> | 0.80 | 0.87 | 0.85 | 0.88 |
|  | 100<sup>th</sup>- 95<sup>th</sup> | 0.78 | 0.86 | 0.81 | 0.86 |
|  | 100<sup>th</sup>- 90<sup>th</sup> | 0.78 | 0.84 | 0.79 | 0.85 |
|  | 100<sup>th</sup>- 50<sup>th</sup> | 0.71 | 0.74 | 0.73 | 0.72 |
|  | 99<sup>th</sup> - 95<sup>th</sup> | 0.95 | 0.96 | 0.99 | 0.99 |
|  | 99<sup>th</sup>- 90<sup>th</sup> | 0.94 | 0.94 | 0.98 | 0.99 |
|  | 99<sup>th</sup>- 50<sup>th</sup> | 0.87 | 0.85 | 0.94 | 0.94 |
|  | 95<sup>th</sup>- 90<sup>th</sup> | 0.99 | 0.98 | 1.00 | 1.00 |
|  | 95<sup>th</sup>- 50<sup>th</sup> | 0.89 | 0.87 | 0.96 | 0.95 |
|  | 90<sup>th</sup>- 50<sup>th</sup> | 0.90 | 0.88 | 0.97 | 0.96 |
| Nodal Averaged <> Element | 100<sup>th</sup> | 0.92 | 0.91 | 0.96 | 0.97 |
|  | 99<sup>th</sup> | 0.99 | 0.96 | 1.00 | 0.99 |
|  | 95<sup>th</sup> | 1.00 | 0.99 | 1.00 | 1.00 |
|  | 90<sup>th</sup> | 1.00 | 0.98 | 1.00 | 1.00 |
|  | 50<sup>th</sup> | 1.00 | 0.90 | 1.00 | 1.00 |

## Discussion

This study evaluated the injury risk functions and predictability of mTBI for different strain post-processing methods and showed that the strain value could differ significantly. A significant difference could be seen for both models when presenting the 100th percentile element value compared to the lower percentile values. For this method, the potential problem with numerical instability can be most pronounced. The lower percentile values and the nodal averaging are two ways of overcoming numerical instability.

A difference in how the choice of strain post-processing influences the risk curves was seen between the two models. For THUMS, the risk curves of the same percentile values with element strain or nodal averaged element strain were most similar, whereas, for the KTH model, this was only the case for the 50^th^ percentile value. For the other curves, one could see that the element value of a percentile was most similar to the nodal averaged element value of the percentile value above, e.g., 99^th^ percentile element value and 100^th^ percentile value with nodal averaging. However, this was only the case for the whole brain for the KTH model and not for the brainstem. The difference between the risk curves is probably due to the difference in the number of nodes and elements. The KTH model has a coarser mesh of around 4,000 elements for the brain compared to THUMS, with around 118,000 elements. The finer mesh has a more continuous strain field without nodal averaging, and this may explain the small difference in risk curves between element strain and nodal averaged element strain for the THUMS model.

The ROC curves and AUC values were analyzed to check the injury predictability. For both models, the 50^th^ percentile values had the highest AUC for the whole brain, but there were minor differences between the different percentile values. There were also minor differences in AUC values between the element and the nodal averaged element values for the same percentile value. These results show that for the current dataset and these two models, the percentile value and how the strain is processed had a minor influence on the injury predictability.

Previous studies have advocated using a lower percentile value than the maximum value with the motivation to remove numerical instability. Especially for the cases with the highest peak strain, one could identify a few elements that had higher element strains than the rest. When taking a percentile value below the 100^th^ percentile value, numerical instabilities can be removed, but also some of the information is lost. One example is the information of the location of the high strain. This is more pronounced when evaluating the whole brain than subregions of the brain since they are smaller, and the location can differ much less. Some previous studies (Anderson et al., 2020; Wu et al., 2020) have suggested that peak strain at subregions is a better predictor for brain injuries. However, this was not the case for the current study when comparing the whole brain and the brainstem, where there was a higher AUC value for the entire brain than the brainstem. In this study, we chose to evaluate the brainstem as a subregion because this was the only subregion that was predefined for both models. In the study by Giordano & Kleiven (2014), the brainstem was the region with the highest AUC for MPS.

In addition to evaluating the injury prediction, the ranking of the simulations was also considered since FE models are used to rate and rank different safety designs. In this study, the severity of the impacts was ranked. For both models, the correlation in the ranking was relatively high (0.71-1.00) when comparing the different outputs. So, the choice of percentile value and element or nodal averaged element value had a small effect on the ranking of the severity.

The linear correlation between the different percentiles and element or nodal averaged element value mainly showed a high correlation. The most significant deviation was found for the cases with higher peak strain (Figure A1 to Figure A4).

The dataset presented by Sanchez et al. (2019) was used in this study. The data is based on impacts in American football, which has a duration of up to 100 ms. A limitation with the dataset used in this study is that some of the impacts have a shorter duration than 100 ms because of various problems with the experimental test. Another limitation with the dataset is the relatively small sample size (20 injured and 33 uninjured). Earlier publications (Anderson et al., 2020; Wu et al., 2020) have used leave-one-out cross-validation to adjust for the small dataset when developing the injury risk functions. In this study, we chose to only create one risk curve for all data points due to the objective of this study.

The focus of this study has been on the first principal Green-Lagrange peak strain with different percentile values and either element value or nodal averaged element value. The Green-Lagrange strain seems to be the most common output used among different FE head models (Fahlstedt et al., 2021). Most models use the peak with different percentile values, but there are also suggestions about other metrics (Bandak and Eppinger, 1995; Sullivan et al., 2015), which has not been evaluated in this study. The focus has also been on the maximum principal strain, whereas recent studies have proposed other alternative strain measures as a better predictor for mTBI (Giordano and Kleiven, 2014; Hajiaghamemar et al., 2020; Ji et al., 2015; Wu et al., 2019; Zhou et al., 2021).

With the results presented in this study, it is important to stress that the same post-processed strain should be used as the one used to develop the injury risk function. The results from this study show that 100^th^ percentile strain with element value could give high strain due to only a few elements in a model. This could be solved by using a strain that is calculated with nodal averaging and/or lower percentile value, but high local strain levels in a few elements due to stability issues should ideally be corrected during model generation.

## Acknowledgement

The authors would like to acknowledge Chris Withnall from Biokinetics and Associates Ltd. for providing the NFL data. MF and SK have partly been financed by FFI (Strategic Vehicle Research and Innovation).

## References

Anderson, E.D., Giudice, J.S., Wu, T., Panzer, M.B., Meaney, D.F., 2020. Predicting Concussion Outcome by Integrating Finite Element Modeling and Network Analysis. Front. Bioeng. Biotechnol. 8.

Atsumi, N., Nakahira, Y., Iwamoto, M., 2016. Development and Validation of a Head/Brain FE Model and Investigation of Influential Factor on the Brain Response during Head Impact. Int. J. Veh. Saf. 9, 1–23.

Bandak, F.A., Eppinger, R.H., 1995. A three-dimensional finite element analysis of the human brain under combined rotational and translational acceleration. Stapp Car Crash J. 38, 145–163.

Clark, J.M., Taylor, K., Post, A., Hoshizaki, T.B., Gilchrist, M.D., 2018. Comparison of Ice Hockey Goaltender Helmets for Concussion Type Impacts. Ann. Biomed. Eng. 46, 986–1000.

Elkin, B.S., Gabler, L.F., Panzer, M.B., Siegmund, G.P., 2019. Brain Tissue Strains Vary with Head Impact Location: A Possible Explanation for Increased Concussion Risk in Struck versus Striking Football Players. Clin. Biomech. 64, 49–57.

Fahlstedt, M., Abayazid, F., Panzer, M.B., Trotta, A., Zhao, W., Ghajari, M., Gilchrist5, M.D., Ji, S., Kleiven, S., Li, X., Annaidh, A.N., Halldin, P., 2021. Ranking and Rating Bicycle Helmet Safety Performance in Oblique Impacts Using Eight Different Brain Injury Models. Ann. Biomed. Eng.

Gabler, L.F., Crandall, J.R., Panzer, M.B., 2016. Assessment of Kinematic Brain Injury Metrics for Predicting Strain Responses in Diverse Automotive Impact Conditions. Ann. Biomed. Eng. 44, 3705–3718.

Gabler, L.F., Crandall, J.R., Panzer, M.B., 2018. Development of a Metric for Predicting Brain Strain Responses Using Head Kinematics. Ann. Biomed. Eng. 46, 972–985.

Ghajari, M., Hellyer, P.J., Sharp, D.J., 2017. Computational Modelling of Traumatic Brain Injury Predicts the Location of Chronic Traumatic Encephalopathy Pathology. Brain 140, 333–343.

Giordano, C., Kleiven, S., 2014. Evaluation of Axonal Strain as a Predictor for Mild Traumatic Brain Injuries Using Finite Element Modeling. Stapp Car Crash J. 58.

Giudice, J.S., Park, G., Kong, K., Bailey, A., Kent, R., Panzer, M.B., 2019. Development of Open-Source Dummy and Impactor Models for the Assessment of American Football Helmet Finite Element Models. Ann. Biomed. Eng. 47, 464–474.

Hajiaghamemar, M., Seidi, M., Margulies, S.S., 2020. Head Rotational Kinematics, Tissue Deformations, and Their Relationships to the Acute Traumatic Axonal Injury. J. Biomech. Eng. 142, 1–13.

Hernandez, F., Giordano, C., Goubran, M., Parivash, S., Grant, G., Zeineh, M., Camarillo, D., 2019. Lateral impacts correlate with falx cerebri displacement and corpus callosum trauma in sports-related concussions. Biomech. Model. Mechanobiol. 18, 631–649.

Ji, S., Zhao, W., Ford, J.C., Beckwith, J.G., Bolander, R.P., Greenwald, R.M., Flashman, L. a, Paulsen, K.D., McAllister, T.W., 2015. Group-Wise Evaluation and Comparison of White Matter Fiber Strain and Maximum Principal Strain in Sports-Related Concussion. J. Neurotrauma 32, 441–454.

Kleiven, S., 2007. Predictors for Traumatic Brain Injuries Evaluated through Accident Reconstructions. Stapp Car Crash J. 51, 81–114.

Newman, J., Beusenberg, M., Fournier, E., Shewchenko, N., King, A., Yang, K., Zhang, L., McElhaney, J., Thibault, L., Gerry McGinnis, 1999. A New Biomechanical Assessment of Mild Traumatic Brain Injury Part I - Methodology. In: Proceedings of the International Research Council on Biomechanics of Injury (IRCOBI) Conference. Sitges, Spain, pp. 17–36.

Newman, J.A., Beusenberg, M.C., Shewchenko, N., Withnall, C., Fournier, E., 2005. Verification of Biomechanical Methods Employed in a Comprehensive Study of Mild Traumatic Brain Injury and the Effectiveness of American Football Helmets. J. Biomech. 38, 1469–81.

Panzer, M.B., Myers, B.S., Capehart, B.P., Bass, C.R., 2012. Development of a Finite Element Model for Blast Brain Injury and the Effects of CSF Cavitation. Ann. Biomed. Eng.

Pellman, E.J., Viano, D.C., Tucker, A.M., Casson, I.R., Waeckerle, J.F., 2003. Concussion in professional football: Reconstruction of game impacts and injuries. Neurosurgery 53, 799–814.

Roozenbeek, B., Maas, A.I.R., Menon, D.K., 2013. Changing Patterns in the Epidemiology of Traumatic Brain Injury. Nat. Rev. Neurol. 9, 231–6.

Sanchez, E.J., Gabler, L.F., Good, A.B., Funk, J.R., Crandall, J.R., Panzer, M.B., 2019. A Reanalysis of Football Impact Reconstructions for Head Kinematics and Finite Element Modeling. Clin. Biomech. 64, 82–89.

Sullivan, S., Eucker, S.A., Gabrieli, D., Bradfield, C., Coats, B., Maltese, M.R., Lee, J., Smith, C., Margulies, S.S., 2015. White matter tract oriented deformation predicts traumatic axonal brain injury and reveals rotational direction-specific vulnerabilities. Biomechancial Model Mechanobiol. 14, 877–896.

Trotta, A., Clark, J.M., McGoldrick, A., Gilchrist, M.D., Annaidh, A.N., 2020. Biofidelic finite element modelling of brain trauma: Importance of the scalp in simulating head impact. Int. J. Mech. Sci. 173, 105448.

Wu, S., Zhao, W., Rowson, B., Rowson, S., Ji, S., 2020. A Network-Based Response Feature Matrix as a Brain Injury Metric. Biomech. Model. Mechanobiol. 19, 927–942.

Wu, T., Alshareef, A., Giudice, J.S., Panzer, M.B., 2019. Explicit Modeling of White Matter Axonal Fiber Tracts in a Finite Element Brain Model. Ann. Biomed. Eng. 47, 1908–1922.

Zhou, Z., Li, X., Liu, Y., Fahlstedt, M., Georgiadis, M., Raymond, S.J., Grant, G., Kleiven, S., Camarillo, D., Engineering, N., Royal, K.T.H., 2021. Towards a comprehensive delineation of white matter tract-related deformation Department of Radiology, Stanford University, Stanford, CA, 94305, USA. Department of Neurosurgery, Stanford University, Stanford, CA, 94305, USA. Department of Neur. bioRxiv 1–33.

