## Supplementary Material for "Influence of Strain Post-Processing on Brain Injury Prediction"

### Appendix

*Table A1. Summary of the two models*

| Brain Model | KTH | THUMS |
|---|---|---|
| **Number of brain elements** | 4.1k | 118.8k |
| **Number of brain nodes** | 5.2k | 128.2k |
| **Number of brainstem elements** | 60 | 1682 |
| **Number of brainstem nodes** | 124 | 2197 |
| **Intracrainal volume [$dm^3$]** | 1.4 | 1.6 |
| **Element Formulation** | Eight-node brick element with selectively reduced integration | Eight-node brick element with constant stress |
| **Brain material properties** | Hyperelastic and vicoelastic (Ogden model with Prony series viscoelasticity) | Linear viscoelastic (standard linear solid) |
| **Interactions between different parts of the brain and the brain-skull interface** | Tangential sliding without separation in normal direction between subdural CSF and brain & between the CSF located on both sides of the falx/tentorium and the brain | Automatic surface to surface between pia sagittal – falx, & brain-tentorium. Continuous mesh between brain and subdural CSF |

*Table A2. Mesh quality of the brain tissue for the two models*

| | THUMS | | | KTH | | |
|---|---|---|---|---|---|---|
| | **Number of failed elements** | **Percentage of failed elements** | **Max/Min** | **Number of failed elements** | **Percentage of failed elements** | **Max/Min** |
| **Warpage < 50°** | 8 | 0% | 53.1 | 4 | 0% | 74.1 |
| **Aspect ratio <8** | 28 | 0% | 8.5 | 6 | 0% | 11.0 |
| **Skew <70°** | 186 | 0% | 77.6 | 0 | 0% | 55.0 |
| **Quad face >30°** | 1054 | 1% | 11.4 | 0 | 0% | 30.5 |
| **Quad face <150°** | 1504 | 1% | 175.5 | 0 | 0% | 149.9 |
| **Normalized Jacobian >0.4** | 0 | 0% | 0.33 | 0 | 0% | 0.32 |

*Table A3. Constants for theWeibull survival analysis and the 50% risk of mTBI (p(50)).*

| | | | | α | β | p50 |
|---|---|---|---|---|---|---|
| KTH | Brain | Element | 100th | 0.470 | 2.527 | 0.407 |
| | | | 99th | 0.313 | 3.746 | 0.284 |
| | | | 95th | 0.252 | 4.065 | 0.230 |
| | | | 90th | 0.224 | 4.170 | 0.205 |
| | | | 50th | 0.124 | 4.986 | 0.116 |
| | | Nodal Averaged Element | 100th | 0.332 | 3.161 | 0.296 |
| | | | 99th | 0.262 | 3.932 | 0.238 |
| | | | 95th | 0.225 | 4.089 | 0.206 |
| | | | 90th | 0.205 | 4.171 | 0.187 |
| | | | 50th | 0.122 | 4.973 | 0.113 |
| | Brainstem | Element | 100th | 0.226 | 1.888 | 0.186 |
| | | | 99th | 0.224 | 1.906 | 0.185 |
| | | | 95th | 0.192 | 1.958 | 0.159 |
| | | | 90th | 0.141 | 2.171 | 0.119 |
| | | | 50th | 0.064 | 4.050 | 0.058 |
| | | Nodal Averaged Element | 100th | 0.171 | 1.981 | 0.142 |
| | | | 99th | 0.170 | 1.982 | 0.141 |
| | | | 95th | 0.152 | 2.005 | 0.126 |
| | | | 90th | 0.133 | 2.020 | 0.111 |
| | | | 50th | 0.062 | 3.999 | 0.057 |
| THUMS | Brain | Element | 100th | 1.817 | 1.674 | 1.459 |
| | | | 99th | 0.545 | 2.957 | 0.481 |
| | | | 95th | 0.384 | 3.292 | 0.343 |
| | | | 90th | 0.328 | 3.411 | 0.295 |
| | | | 50th | 0.184 | 4.089 | 0.168 |
| | | Nodal Averaged Element | 100th | 1.229 | 1.816 | 1.004 |
| | | | 99th | 0.523 | 2.944 | 0.462 |
| | | | 95th | 0.376 | 3.258 | 0.336 |
| | | | 90th | 0.324 | 3.389 | 0.290 |
| | | | 50th | 0.183 | 4.067 | 0.168 |
| | Brainstem | Element | 100th | 0.647 | 1.135 | 0.468 |
| | | | 99th | 0.378 | 1.467 | 0.294 |
| | | | 95th | 0.150 | 2.078 | 0.126 |
| | | | 90th | 0.104 | 2.407 | 0.090 |
| | | | 50th | 0.055 | 1.599 | 0.044 |
| | | Nodal Averaged Element | 100th | 0.573 | 1.158 | 0.418 |
| | | | 99th | 0.330 | 1.290 | 0.249 |
| | | | 95th | 0.128 | 2.113 | 0.107 |
| | | | 90th | 0.093 | 2.432 | 0.080 |
| | | | 50th | 0.056 | 1.452 | 0.043 |

The equation of the Weibull risk curve:

$$p = 1 - e^{-\left(\frac{x}{\alpha}\right)^{\beta}}$$

where x is strain. The constants α and β are found in Table A3.

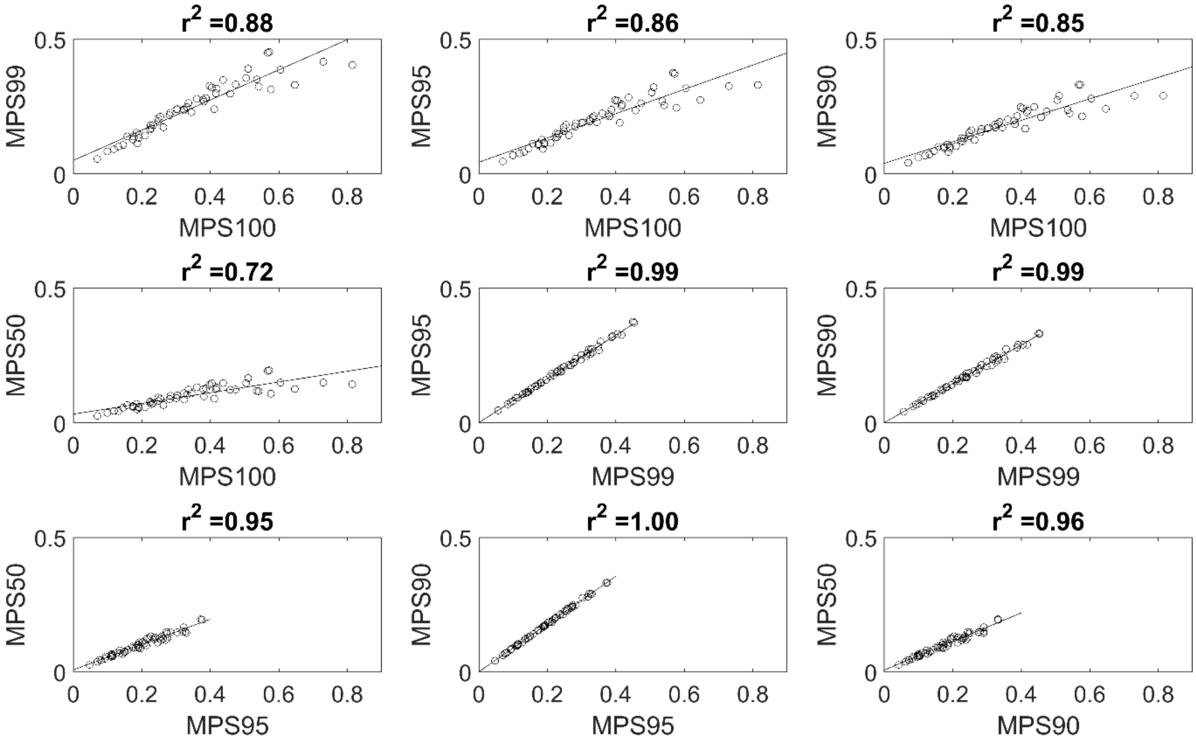


*Figure A1. KTH element value*

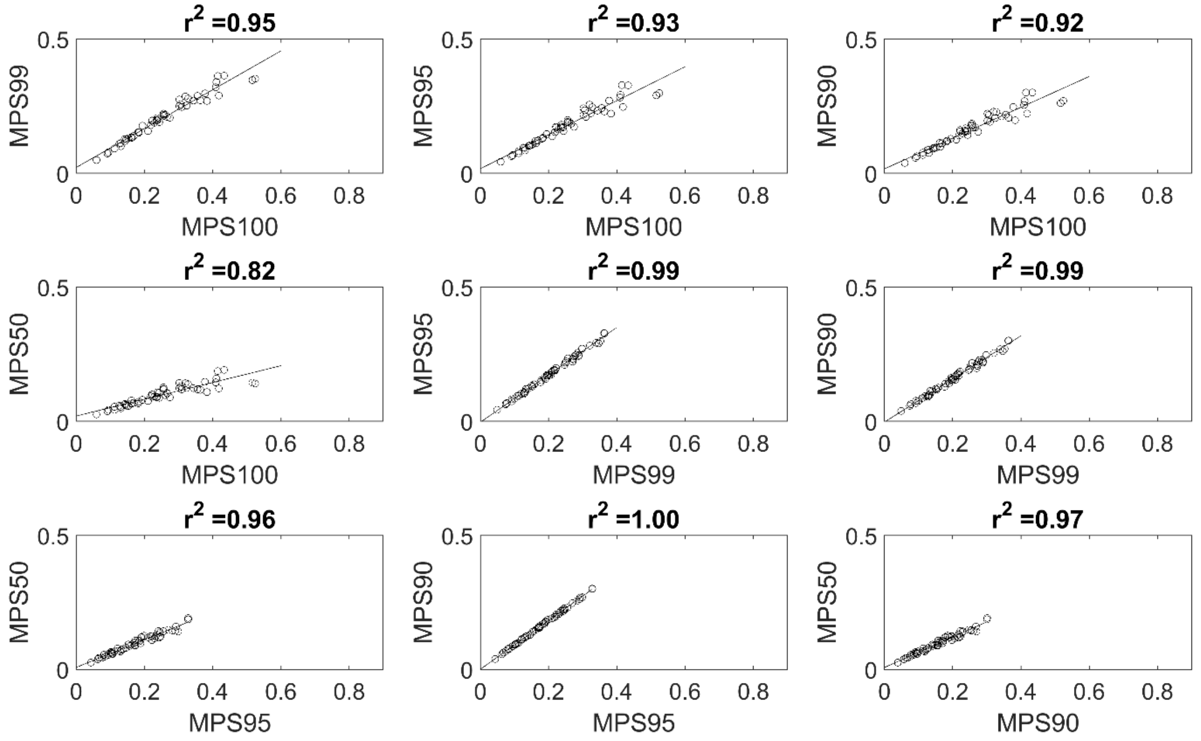


*Figure A2. KTH nodal averaged element value*

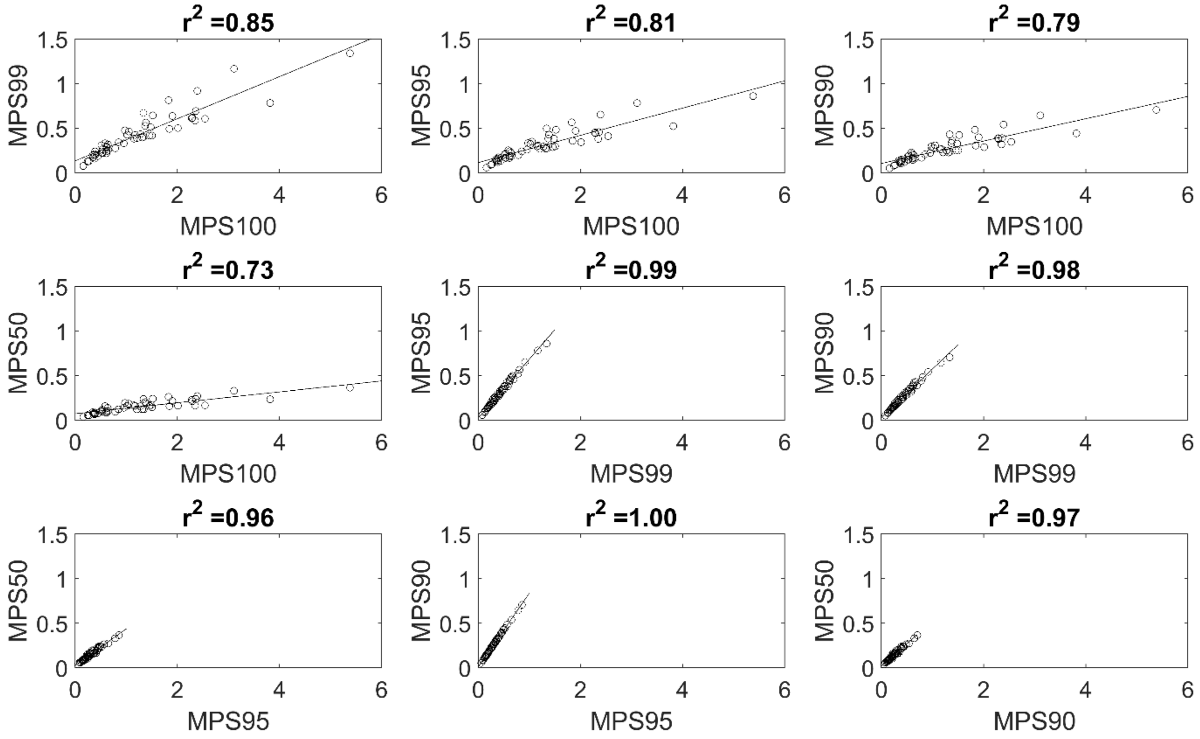


*Figure A3. THUMS element value*

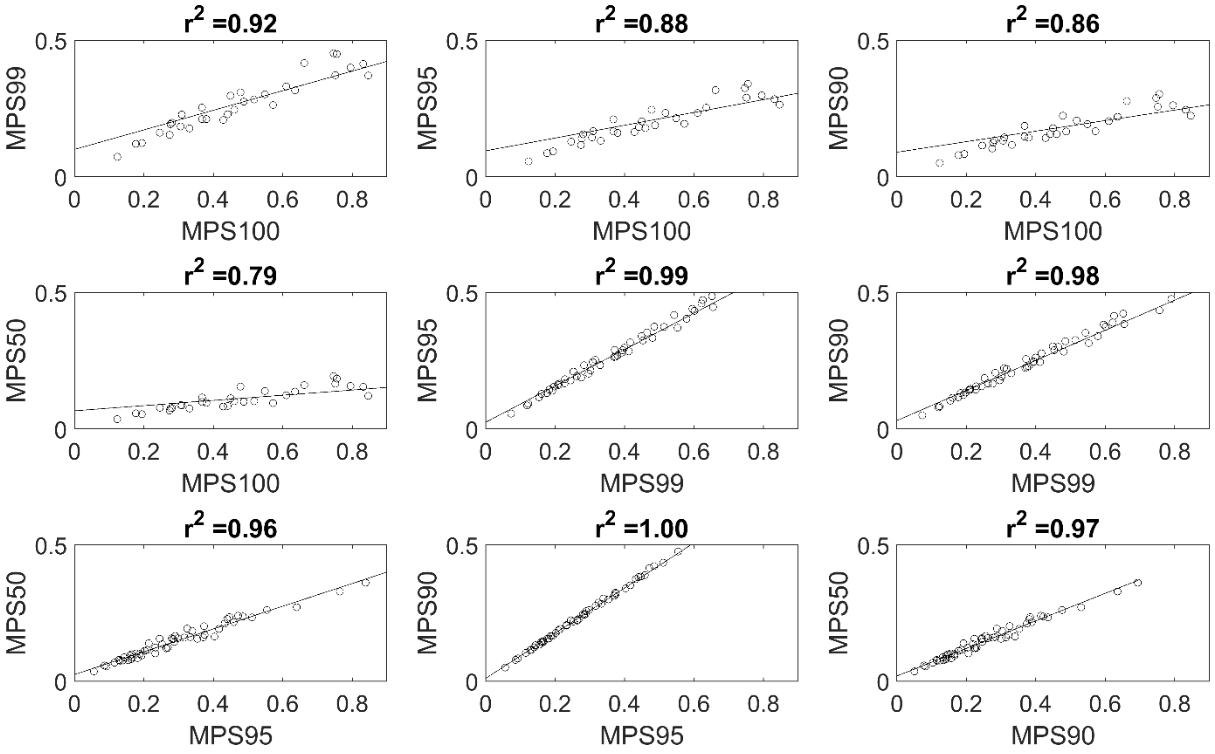


*Figure A4. THUMS nodal averaged element value*
